# A genome-scale CRISPR deletion screen in Chinese Hamster Ovary cells reveals essential regions of the coding and non-coding genome

**DOI:** 10.1101/2025.04.28.650989

**Authors:** Federico De Marco, Ivy Rose Sebastian, Antonino Napoleone, Alexander Molin, Markus Riedl, Nina Bydlinski, Krishna Motheramgari, Mohamed K. Hussein, Lovro Kramer, Thomas Kelly, Thomas Jostock, Nicole Borth

**Author notes:** Nicole Borth, Department of Biotechnology, University of Natural Resources and Life Sciences (BOKU), Vienna, Muthgasse 18, 1190 Vienna, Austria. These authors contributed equally (Federico De Marco, Ivy Rose Sebastian, Antonino Napoleone).

## Abstract

The biopharmaceutical sector relies on CHO cells to investigate biological processes and as the preferred host for production of biotherapeutics. Simultaneously, advancements in CHO cell genome assembly have provided insights for developing sophisticated genetic engineering strategies. While the majority of these efforts have focused on coding genes, with some interest in transcribed non-coding RNAs (e.g., microRNAs and lncRNAs), there remains a lack of genome-wide systematic studies that precisely examine the remaining 90% of the genome. This unannotated “dark matter” includes regulatory elements and other, poorly understood or characterized functionality of the genome that may be potentially critical for cell survival. In this study, we deployed a genome-scale CRISPR screening platform with 112,272 paired guide RNAs targeting 14,034 genomic regions for complete deletion of 150 kb long sections. This platform enabled the execution of a negative screen that selectively identified dying cells to determine regions essential for cell survival. By using paired gRNAs, we overcame the intrinsic limitations of traditional frameshift strategies, which will likely have little or no effect on the non-coding genome. This study revealed 427 regions essential for CHO survival, many of which currently lack gene annotation or known function. For these regions we present annotation status, transcriptional activity as well as annotated chromatin states such as enhancers. Selected regions, specifically those that were negative for all the above, were individually deleted for confirmation. This work sheds a novel light on a substantial part of the mammalian genome which is traditionally difficult to investigate and therefore, neglected.

## 1 Introduction

Advances in genomics are rooted in prior knowledge and the last two decades have gathered an enormous encyclopedia of information with the promise of shaping our current understanding of genome functionality. Multiple projects have yielded for the first time (nearly) complete genome sequences for many scientifically relevant mammalian organisms (Mouse Genome Sequencing Consortium, 2002; Rat Genome Sequencing Project Consortium et al., 2004; Rupp et al., 2018; Venter et al., 2001). However, alongside this wealth of genome sequence information, there has always been a pressing need for large-scale, systematic approaches to sift through it and to understand its respective functionality. With the advent of CRISPR-mediated techniques (Cong et al., 2013; Jinek et al., 2012), there has been a significant expansion in the repertoire of available genetic screening tools. In this context, pooled screens, where each individual cell undergoes its own specific genetic perturbation, have proven to be an effective means of characterizing and correlating the genotype with the phenotype (Shalem et al., 2014; T. Wang et al., 2014). The main principle of pooled screens is that phenotypic selection leads to an enrichment or depletion of genetic perturbations relevant to the phenotype (Bock et al., 2022) allowing for the exploration of thousands of perturbations in parallel (Sanjana, 2017). In general, while the role of CRISPR screens was immediately central to functional genomics in human cells (Konermann et al., 2015; Zhou et al., 2014), it also holds significant potential for many other mammalian cells, extending beyond basic research to include industrially relevant applications.

Mammalian, in particular Chinese hamster ovary cells (CHO) are the most used expression system for the production of recombinant proteins (Fischer et al., 2015; J. Y. Kim et al., 2012; Lai et al., 2013), where CHO cells produce 95 of 107 recombinant products on the market (Walsh & Walsh, 2022). This popularity is largely due to the ability of this production platform to grow to high cell densities in suspension, to produce high yields of complex recombinant proteins with human-like post-translational modifications, and to exhibit restricted susceptibility to viral infections (Berting et al., 2010; Lewis et al., 2013). Since their inception in 1987 as the production platform for recombinant tissue plasminogen activators (Griffiths & Electricwala, 1987), CHO cell lines have played a dominant role in the biopharmaceutical sector. In 2018, a more accurate and complete genome of *Cricetulus griseus*, the Chinese hamster, became available (Rupp et al., 2018). Since then, the Chinese hamster community has made further advancements, resulting in a comprehensive set of stable reference chromosome coordinates (Hilliard et al., 2020) that can be used to enhance our understanding of the genomic mechanisms, dynamics, and evolution of this widely used biopharmaceutical cell line. Additionally, the industrial relevance of CHO cells should not overshadow their long-lasting role as a model system for elucidating fundamental scientific principles concerning the functionality of many coding genes (Maeda et al., 2006; Rolig et al., 1997; Urlaub et al., 1983).

However, as for many other mammalian organisms, a considerable portion of the CHO genome remains uncharacterized or less explored. Typical genome-wide screens primarily focus on coding genes (Karottki et al., 2021; S. H. Kim et al., 2023; Xiong et al., 2021), which account for only 3% of the total genetic information (Lewis et al., 2013), with reduced attention given to transcribed but non-coding RNAs, such as microRNAs (Diendorfer et al., 2015; Maccani et al., 2014; Raab et al., 2019) and long non-coding RNAs (Motheramgari et al., 2020; Novak et al., 2023; Schmieder et al., 2021; Wright & Sanjana, 2016). Nevertheless, there is a lack of systematic, genome-wide studies to comprehensively investigate the remaining 90% of the genome. This “dark matter” potentially includes regulatory elements and poorly annotated genomic regions, potentially involved in structural functions that affect nuclear architecture and chromosome territories. The concurrent availability of a well annotated and assembled genome (Hilliard et al., 2020; Rupp et al., 2018) coupled with the accessibility of advanced CRISPR strategies that enable full deletion of genome regions, rather than the traditional frameshift knockouts applicable only to coding genes (Schmieder et al., 2018), now enables genome-wide studies in CHO cells, allowing to study relevant properties across both the coding and non-coding genome. To date, no attempt has been made to conduct a truly genome-wide screen that covers 100% of the genome in its entirety.

In this study, we deployed a genome-wide CRISPR-Cas9 deletion screen with 112,272 paired guide RNAs targeting 14,034 genomic regions for systematic large-scale deletions of 150 kb DNA fragments. With this platform, we performed a negative screen that actively identified dying cells that had lost genome regions essential for survival. By utilizing paired gRNAs, we overcame the intrinsic limitations of a traditional frameshift screen, which would likely have minimal or unknown effects on the non-coding genome (Canver et al., 2014). Our work revealed 427 regions essential for CHO cell survival, for which we present data on their annotation (or lack thereof), transcription (if any), and annotated chromatin states. Selected regions, particularly those with no annotation, were individually deleted for confirmation. Our study sheds novel light on a substantial portion of the mammalian genome that has traditionally been difficult to investigate and, as a result, often neglected. These findings represent a significant milestone, enabling novel approaches to understanding the functional roles of what has long been erroneously called “junk DNA”.

## 2 Materials and methods

### 2.1 Cell cultures

Suspension-adapted CHO K1 cells (ECACC, 85051005) were cultured in CD-CHO medium (Thermo Fisher Scientific, #10743029) supplemented with 8 mM L-glutamine (Sigma-Aldrich, #G7513) and 2 μL/mL Anti-Clumping Agent (Thermo Fisher Scientific, #0010057AE) at 37 °C, 7% CO_2_, in a shaking incubator. Cells were initially grown in TPP Tube-Spin Bioreactors (TPP, #TPP-87050) and scaled up into shake flasks (Corning, #431143) for upscaling experiments. CHO K1 cells were passaged every 3-4 days, and cell viability and viable cell density (VCD) were monitored using the Vi-CELL XR Cell Viability Analyzer (Beckman Coulter). All cell lines used in this study were confirmed negative for mycoplasma contamination.

### 2.2 Cloning and plasmid construction

All the plasmids and oligonucleotide sequences used in this study are listed in **Suppl. table S1A, S1B**. Parental plasmids pSQT1313 and pLL3.7 (Addgene, Inc.) were modified for pgRNA cassette insertion via BsmBI restriction sites by first cloning the hU6 promoter and one of the two modified SpCas9 gRNA scaffolds into the base vectors via restriction enzyme digestion (XbaI/XhoI, NEB) and ligation (denoted as pSQT1313_GB1 and pLL3.7_GB1, respectively). The lentiviral plasmid, pLL3.7_GB1, was further modified via Gibson assembly to insert a T2A_BSD sequence to allow for blasticidin selection after transduction. Final pgRNA expression plasmids were constructed through a two-step protocol (**Suppl. figure 3**): insertion of the pgRNA sequences via Gibson cloning, followed by insertion of the 7SK promoter and second SpCas9 gRNA scaffold using BsmBI-mediated (NEB, #R0739S) Golden Gate cloning. Constructs were verified by Sanger sequencing (Eurofins Genomics) and purified with the GeneJet Endo-Free Plasmid Maxiprep Kit (Thermo Scientific, #K0861).

### 2.3 Computational design of pgRNAs

The design of spCas9 pgRNAs was similarly designed as reported in Schmieder et al., 2021 (Schmieder et al., 2021). Briefly, the 2 kb flanking region upstream and downstream of each of the five targeted genes was evaluated for possible PAM sequences using CRISPOR (https://crispor.gi.ucsc.edu/). Instead of calculating the CFD score, the guides output was refined by incorporating the Doench scoring system (Doench et al., 2016) as best suited for CRISPR screens in the context of mammalian genomes. Filtered guides were then mapped against the whole genome (Hilliard et al., 2020) to identify off-targets using the Bowtie alignment software (v1.2.2). Alignment is performed with options to prioritize finding all possible alignments over speed (-h) and report all alignments per each guide (-a) without allowing any mismatch (-n=0) in a seed length of 12 nt (-l=12). As the alignment tool requires the seed to be at the beginning of the sequence, reverse compliment sequences of the guides are mapped and later corrected for orientation while parsing the BAM file. However, to avoid non-unique alignments which are possible with mismatches outside the seed sequence, only guides with zero mismatches are selected for further use. Finally, the guides with any non-unique alignment are also filtered out. For each target gene, the top five scoring pgRNA sequences were then reported (**Suppl. table S1D**).

### 2.4 Verification of pooled pgRNA deletion functionality

pgRNA sequences targeting five non-essential genes (**Suppl. table S1D**) were synthesized as lyophilized 138 bp Ultramer™ DNA oligonucleotides (IDT, Inc). These oligonucleotides were resuspended to 100 µM stock solutions, as per the manufacturer’s instructions. Individual oligomers were amplified by PCR and cloned using the two-step cloning protocol described above, with rank 1 and rank 5 guides pooled separately into mini pools (designated as P1 and P5, respectively) in equimolar ratios. Following the generation of the mini-pool plasmid libraries, 6 x 10^6^ viable cells were co-transfected with the SpCas9 vector (Addgene #129727) at a 1:1 ratio for a total of 24 µg using nucleofection. A negative control with the empty unmodified parental vector was included. Two days post-transfection, mCherry expression was analyzed via flow cytometry, yielding transfection efficiencies of 70-80%. Four days after transfection, genomic DNA (geDNA) and total RNA were isolated from 4 x 10^6^ viable cells. DNA and RNA concentrations and qualities were measured using a Nanodrop One (Thermo Fisher Scientific). geDNA was isolated using the DNeasy® Blood & Tissue Kit (Qiagen), according to the manufacturer’s protocol, and used for deletion PCR analysis. Deletion PCRs were conducted using the GoTaq G2 DNA polymerase (Promega, #M7848) for a total of 20 µL and a 50 ng DNA input per reaction (95 °C for 2 min; 35 cycles: 95 °C for 30 s, 56/59 °C for 60 s, 72 °C for 90 s; 72 °C for 5 min). Total RNA was isolated using the Quick-RNA Miniprep Kit (Zymo, #R1055). mRNA expression for the target genes were assayed by qRT-PCR as previously described (Marx et al., 2021). Deletion PCR and qPCR primers are listed in **Suppl. table S1C**.

### 2.5 Computational design of custom genome-scale pgRNA library

Custom scripts were used to process and filter the PICRH genome to generate a library of pgRNAs. First, the genome was filtered to remove assembled fragments shorter than 500 kb nucleotides. The BEDtools suite (Quinlan & Hall, 2010) was used to generate 150 kb windows of equal length. The windows were trimmed to remove regions that were within 10 kb of the ends of assembled contigs. Subsets of 1 kb regions at either end of each window were fed into *multicrispr* for prediction of pgRNAs. Prospective guide RNAs were removed if they fell within 50 bp upstream or downstream or within the exons of a gene. pgRNAs were only retained if they possessed a Doench2016 score ≥ to 0.5 and had only 1 target with zero off-targets. For each window, eight pgRNAs, if available, were selected based on their Doench score ranking.

### 2.6 Construction of the genome-scale pgRNA plasmid library

The genome-scale library, comprising 112,272 unique guide RNA pairs targeting 14,034 genomic loci, was custom-synthesized as 138 bp oligomers (Twist Bioscience, San Francisco, CA). Upon receipt, the lyophilized oligo pool was reconstituted in nuclease-free TE buffer (pH 8.0) to a final concentration of 10-20 ng/μL. Library amplification was performed in a pooled format via low-cycle PCR using KAPA HiFi HotStart ReadyMix (Roche, #KK2602) according to the manufacturer’s protocol. The amplified PCR products were subsequently integrated into the BsmBI-digested modified lentiviral vector (pLL3.7_GB1_T2A_BSD) using the commercial Gibson assembly kit (NEB #E2611L). The reaction was scaled up 5-fold, with 100 ng of backbone DNA at a 1:5 vector: insert molar ratio. Purified assembly products were electroporated into MegaX DH10B T1R cells (Thermo Scientific, #C640003) to generate intermediate plasmids. These constructs were then subjected to BsmBI digestion and ligated to the SpCas9 scaffold-7SK promoter fragment using T4 ligase. The resulting ligation mixture was transformed into fresh MegaX DH10B T1R cells to produce the final genome-scale plasmid library. Quality control was performed by double-digesting an aliquot of the plasmid library with NdeI and NspI restriction enzymes (NEB). The resulting 233 bp fragment, containing the first gRNA of each pair, was isolated from a 2% agarose gel using the Hi Yield Gel/PCR DNA Fragment Extraction Kit (SLG, #HYDF100). The purified DNA was then subjected to Illumina sequencing to assess library coverage and guide representation.

### 2.7 Lentiviral production and transduction of pooled library

Lentiviral production and transduction were performed as described in Napoleone et al, 2025. Briefly, HEK293T Lenti-X packaging cells (Takara/Clontech, #632180) were seeded into Falcon 525 cm² 3-layer Multi-Flasks (Corning, #11587421) and Falcon 875 cm² 5-layer Multi-Flasks (Corning, #11597421) at 40% confluency. After 24 hours, the cells were transfected using a third-generation lentiviral system comprising the transfer library plasmid, packaging plasmids pMDLg/pRRe (Addgene #12251) and pRSV-Rev (Addgene #12253), and the CMV-VSVG envelope plasmid (Oxgene #OG592) at a mass ratio of 1.5:2:1:1, respectively. Transfection was carried out using the jetPRIME reagent (Polyplus, #101000001). Viral supernatants were harvested at 48 and 96 hours post transfection, clarified by centrifugation at 1000×g for 10 minutes, and filtered through 0.45 µm low protein-binding filters (Millipore, #S2HVU05RE). The clarified supernatants were concentrated 100-fold using a custom 4X lentiviral concentrator solution based on polyethylene glycol (PEG)-8000 (Promega, #V3011) precipitation at a 1:3 ratio and subsequently stored at −80°C. Viral preparations were titrated, and functional titers were assessed using flow cytometry by measuring the transduction efficiency in positively transduced cells expressing the eGFP marker encoded in the library delivery vector. Target CHO K1 cells were transduced using a two-step static transduction protocol (Napoleone et al., 2025), designed to accommodate the lower susceptibility of this cell line to lentiviral infection. Transduction was aimed at 14-16% efficiency (MOI of 0.16) to minimize the likelihood of multiple integrations. A total of 375 million cells were seeded across ten 145 mm cell culture dishes (Greiner Bio-One #639160) and transduced with the lentiviral library at a target coverage of 500X in the presence of 8 µg/mL polybrene (Merck Millipore, #TR-1003-G). After 24 hours, cells were resuspended in fresh media without virus and transferred to shaking conditions in 1 L flasks (Corning #431147). Following this, positively transduced cells were selected with 10 μg/mL blasticidin (Invivogen #ant-bl-1) for 9 days.

### 2.8 Vector copy number analysis using ddPCR

After antibiotic selection of the transduced cell library, geDNA was extracted using the DNeasy® Blood & Tissue Kit (Qiagen), according to the manufacturer’s protocol. Vector copy number (VCN) of the integrated gRNA was determined by ddPCR using the QX200 Droplet Digital PCR System (Bio-Rad Laboratories, Inc) using the EvaGreen DNA-binding dye (Bio-Rad #1864033) according to the manufacturer’s guidelines, unless stated otherwise. Genomic DNA samples (1 µg) were digested with HindIII. The ddPCR mix contained 1X QX200™ ddPCR™ EvaGreen Supermix, 0.25 µM primers (**Suppl. table S1C**) and 100 ng digested geDNA in a final volume of 22 µL with a mastermix to input DNA ratio of 1:4. Droplets were generated using the QX200™ Droplet Generator and amplified in a C1000 Touch thermal cycler. The amplification was performed with a ramp rate of 2 °C/sec and the following cycling conditions: initial denaturation at 95 °C for 5 min, 40 cycles at 95 °C for 30 s and 60 °C for 1 min, followed by a signal stabilization step at 4 °C for 5 min, and a final step at 90 °C for 5 min. Data acquisition and analysis were performed with the QX200™ Droplet Reader and QuantaSoft™ Software (Bio-Rad Laboratories, Inc), with FUT8 (copy no. = 2) as the reference gene for normalization. The copy number was calculated from the absolute ddPCR quantification value obtained as concentration (copies /µL), using the following formula:

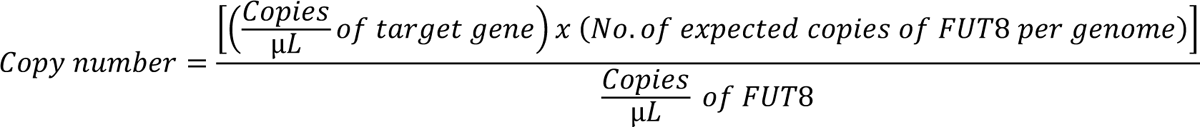

### 2.9 Pooled CRISPR-Cas9 dropout screening procedure

The cell library underwent two rounds of transfection with 30 µg of the SpCas9 plasmid, spaced one week apart. Untransfected pgRNA-containing library cells were used as controls. To ensure a coverage of >500 cells per pgRNA, six replicates of 10 × 10^7^ cells were seeded at a density of 0.5 × 10^6^ cells/mL in 500 mL Erlenmeyer flasks (Thermo Scientific Nalgene, #10686673) containing 200 mL of the standard medium. Cells were passaged every three days for a total of 21 days, after which 90 × 10^6^ cells were harvested for genomic DNA extraction.

### 2.10 PCR retrieval of pgRNA sequences and Illumina sequencing

Total genomic DNA was purified from 3–10 ml of cell suspension. Up to 90 x 10^6^ cells from each replicate were centrifuged at 200 x g for 6 minutes and the pellet was resuspended in 10 ml of sterile PBS. geDNA was isolated using QIAamp DNA Blood Maxi kit (Qiagen, #51194). The original protocol does not include an RNAse treatment step. The presence of RNA contamination can skew absorbance-based quantification methods, and large amounts of RNA can negatively impact subsequent PCR steps. Thus, samples were treated with RNase A (Qiagen, #19101) for 5 minutes prior to the start of the geDNA isolation protocol. Purified geDNA was eluted in Buffer AE (10 mM Tris-Cl; 0.5 mM EDTA; pH 9.0) and the concentrations were quantified using a Qubit fluorometer (Thermo Fisher Scientific). Multiple PCR reactions were prepared to amplify the total harvested geDNA, ensuring a minimum coverage of 500-fold per cell per perturbation. For PCR retrieval of pgRNA sequences, geDNA was divided into 50 μL reactions such that each tube had at most 5 μg geDNA. Each reaction contained 2.5 μL of each primer (10 μM), 4 μL of dNTPs (2.5 mM), 5% DMSO (DeWeirdt et al., 2021), 5 μL of 10X Titanium Taq Buffer, and 0.75 μL of 10X Titanium Taq DNA Polymerase (Takara Bio, #639209). PCR cycling conditions were as follows: an initial denaturation step of 3 minutes at 95 °C; followed by 30 s at 95 °C, 30 s at 62 °C, 30 s at 72 °C, for 28 cycles; and a final 10 minute extension step at 72 °C. PCR primers, pgRNA_gRNA1_fw (5’-TCTTGTGGAAAGGACGAAACAC) and pgRNA_gRNA1_rv (5’-CGCCGATGAATAGCGTGAGA), were synthesized by Integrated DNA Technologies. Samples were purified with Agencourt AMPure XP Beads according to the manufacturer’s instructions (Beckman Coulter, #A63881). The resulting amplicons (162 bp) were sequenced on an Illumina NovaSeq S4 using 150 bp x 2 paired-end sequencing (Vienna BioCenter NGS facility).

### 2.11 Computational analysis of screen data

The raw sequencing data were obtained from paired-end, non-directional Illumina sequencing. Two distinct methods were employed to generate the read count tables. In the first method, the raw data in FASTA format were processed using *‘mageck count’* (Li et al., 2014) to generate a straightforward gRNA count table leveraging its default settings. The second method involved several preprocessing steps. First, raw paired-end reads were merged using PEAR (Paired-End reAd merger) (Zhang et al., 2014). Then, the merged reads were corrected for orientation and trimmed to retain only the pgRNAs based on known flanking sequences, accepting sequence lengths of ∼20 bp (18<pgRNA<22). The alignment to the reference library was performed using Bowtie2 (Langmead et al., 2019) with settings *“--norc -n 0 --end-to-end”* (no reverse complement, maximum mismatch in seed = 0, end-to-end alignment) and converted to .bam files using Samtools (Danecek et al., 2021). These files were then summarized into a count table using *‘mageck count’* with default settings. For downstream analysis, MAGeCK VISPR was used to assess the data from both methods. MAGeCK-VISPR applies a maximum likelihood estimation (MLE) algorithm to evaluate gene essentiality based on gRNA enrichment data. All analysis pipelines are available https://github.com/NBorthLab/CHO-essentialome.

### 2.12 Functional enrichment analysis

Enrichment analysis of Gene Ontology (GO) terms was conducted using over-representation analysis implemented in clusterProfiler v4.10.0 (T. Wu et al., 2021) for R v4.3.3. The functional annotation based on the RefSeq genome of *Cricetulus griseus* (GCF_003668045.3) was retrieved using AnnotationHub v3.10.0 (accession ‘AH114610’, snapshot date ‘2023-10-21’). The over-representation analysis used a one-sided Fisher’s exact test for hypothesis testing and multiple testing correction by Benjamini-Hochberg (Benjamini & Hochberg, 1995) with an adjusted *p*-value cutoff of 0.05 to define significantly enriched GO terms. The background used for statistical testing comprised all annotated genes in the Chinese hamster genome. Redundant enriched GO terms were merged using the Wang semantic similarity (J. Z. Wang et al., 2007) measure with a threshold of 0.7.

### 2.13 Assessment of transcription in essential regions

To determine active transcription within essential regions, the RNA-seq dataset from (Weinguny et al., 2020) (BioProject PRJEB37009) was used. In particular, the two samples corresponding to CHO-K1 with supplementation of 8mM ʟ-Glutamine were analyzed. The RNA-seq data was processed and aligned to the *Cricetulus griseus* RefSeq genome as described by Riedl et al. (Riedl et al., 2025). For quantification, genomic regions were segmented into bins of 10bp and used as features in the featureCounts function of Rsubread (v3.20). Multiple feature overlaps were enabled to allow reads to be assigned to several adjacent bins. Read counts were normalized using the trimmed means of M-values method of edgeR (v4, https://doi.org/10.1101/2024.01.21.576131). Normalized log_2_ transformed counts per million (CPM) were averaged between the two RNA-seq samples. Essential regions were considered transcribed if the normalized log_2_CPM exceeded zero in at least nine consecutive bins.

### 2.14 Chromatin states overlap enrichment analysis

Using the PICRH assembly, eleven chromatin states were learned from published ChIP-seq data (Feichtinger et al., 2016). Briefly, patterns of six histone marks were used to train a hidden Markov model with ChromHMM v1.24 for the Chinese hamster RefSeq genome GCF_003668045.3. Time point three (53 hours) was used for overlap enrichment of essential genomic regions. Chromatin state segments of 200bp were only considered overlapping if they fully overlapped with genomic regions. Enrichment of a chromatin state *i* was calculated as

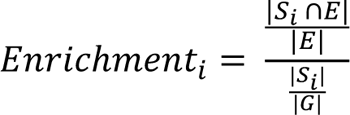

where *S*_*i*_ denotes the segments of chromatin state *i*, *E* the segments overlapping with essential regions, and *G* the segments overlapping with all regions, used as background. Enrichment of chromatin states was further assessed statistically by performing a permutation test using the R package regioneR v1.34.0 (Gel et al., 2016). Regions were resampled from all genomic regions 2000 times. Resulting *p*-values were adjusted for multiple testing using the Benjamini-Hochberg correction.

### 2.15 Individual pgRNA cloning and assay for genomic deletion

The sequences of six depleted pgRNAs from the screen were derived from the 112k library and synthesized as lyophilized 138 bp Ultramer DNA oligonucleotides (IDT, Inc). The guide oligonucleotides for each selected region were individually cloned following the same protocol described in “Verification of pooled pgRNA deletion functionality”. Final plasmid sequences were confirmed by Sanger sequencing (Eurofins Genomics). CHO K1 cells were transiently co-transfected with pgRNAs for a given region and the SpCas9 vector (Addgene #129727) via the Neon Transfection System (Invitrogen). Each group was transfected with a total of 30 µg of DNA per 10 x 10^6^ cells. Transfection efficiency was examined after 24 hours using flow cytometry. Cell viability, total cell numbers and live cell size were collected daily for six days. Confirmation of genomic deletion was assayed via deletion PCR, following the standard protocol previously described. All the primer sequences and primer combinations used for the characterization of the genomic deletions are described in the **Suppl. table S1C**.

### 2.16 Flow cytometric analysis of cell apoptosis

For flow cytometry-based apoptosis analysis, CHO K1 cells were transfected with the combination of SpCas9 and the pgRNAs targeting each essential region. Cells were collected after 24 hours for analysis. A total of 5 × 10⁵ cells were washed with PBS, resuspended in 500 μL of Zombie NIR dye (BioLegend, #423105) at a 1:500 dilution, and incubated at room temperature in the dark. After 15-30 minutes, cells were washed with an additional 1 ml of PBS and incubated with CellEvent Caspase-3/7 Detection Reagent (Thermo Scientific, #C10431) following the manufacturer’s protocol. Briefly, effector caspases such as caspase-3 and caspase-7, are key mediators of apoptosis, and their activation serves as a hallmark of programmed cell death. The percentage of apoptotic cells was characterized by flow cytometry (Cytoflex S, Beckman Coulter), based on the signal for Texas Red (590/610 nm) for the caspase detection reagent and Zombie NIR (633/746 nm) for live/dead cell discrimination. At least 15,000 events were recorded for each sample using the following emission filter settings: 600/BP60 and 665/BP30.

### 2.17 Statistical analysis

All statistical analysis was performed using R version 4.3.3. Hypothesis testing between control sample and its stressed counterpart was performed using the Student’s t-test with Benjamini-Hochberg (BH) adjusted p-values unless stated otherwise. All data are presented as means ± SD. Statistical differences are indicated as follows: ns (not significant) P > = 0.05, *P < 0.05, **P < 0.01, and ***P < 0.001.

## 3 Results

### 3.1 Optimization of an efficient, scalable approach to delete large genomic regions by paired guide RNAs

Functional CRISPR screens are typically performed using pooled libraries of single-guide RNAs (sgRNAs). Here, gene disruption relies entirely on inducing frameshift mutations at specific sites to ensure the loss of function of a coding gene (Chakrabarti et al., 2019; Lieber, 2010). This strategy becomes less effective for targeting the non-coding genome, as the functional effects of frameshifts are not predictable here (Canver et al., 2014). To overcome this limitation, paired guide RNAs (pgRNAs) have been shown to efficiently delete genomic regions up to 150 kb in CHO cells (Schmieder et al., 2018), enabling complete removal of the region of interest, resulting in loss of function however this is encoded. This was shown to enable the knockout of non-coding RNAs such as miRNAs (Raab et al., 2019) or lncRNAs (Novak et al., 2023), but should also be effective for deletion of non-transcription based functionality of the genome. To validate a pgRNA-based approach for high-throughput genomic deletions, and to test a dual-promoter strategy, we selected five genes non-essential to cell survival (Fukuda et al., 2019; Hart et al., 2015; Kol et al., 2020). These genes were chosen to allow for deletions of variable sizes ranging from 15 to 100 kb (**Suppl. figure 1**) and were targeted in a mini-pool deletion assay (**Figure 1a**). For pgRNA expression, with each guide being expressed from an individual promoter, we adopted a dual-promoter system to mitigate recombination within promoter sequences (Gasperini et al., 2017). Until now, only the human U6 Pol III promoter (hU6) has been routinely used for expressing gRNAs in deletion strategies employing paired guides (Zhu et al., 2016). Moreover, no published data was found on the use of other Pol III promoters in CHO cells. Studies in HEK293 cells have demonstrated comparable promoter strength for the Pol III promoters mouse U6 (mU6), 7SK and H1 (Kabadi et al., 2014). Therefore, combinations of hU6/7SK and hU6/mU6 were evaluated for their deletion efficiency in targeting glycosyltransferase genes (**Suppl. figure 2**). The 7SK promoter was selected for its advantages, including its smaller size of 270 bp, and the absence of regions likely to recombine with the hU6 promoter. Additionally, an optimized gRNA scaffold was used with both promoters to increase deletion efficiencies (Dang et al., 2015). To verify our strategy, pgRNAs ranking first and fifth by Doench score were chosen for each of the selected target genes (**Suppl. table S1D**) and cloned into the expression cassette via a two-step cloning protocol (Zhu et al., 2016) (**Suppl. figure 3**). Two mini pools labelled P1 for all guides that ranked 1^st^ and P5 for all guides ranking 5^th^ were generated. Transfection efficiencies were assessed using flow cytometry 24 hours post transfection. Deletion PCR confirmed successful genomic deletions (**Figure 1c**), while qRT-PCR showed reduced expression levels for individual genes in the mini-pools (**Figure 1d**).

**Figure 1.**
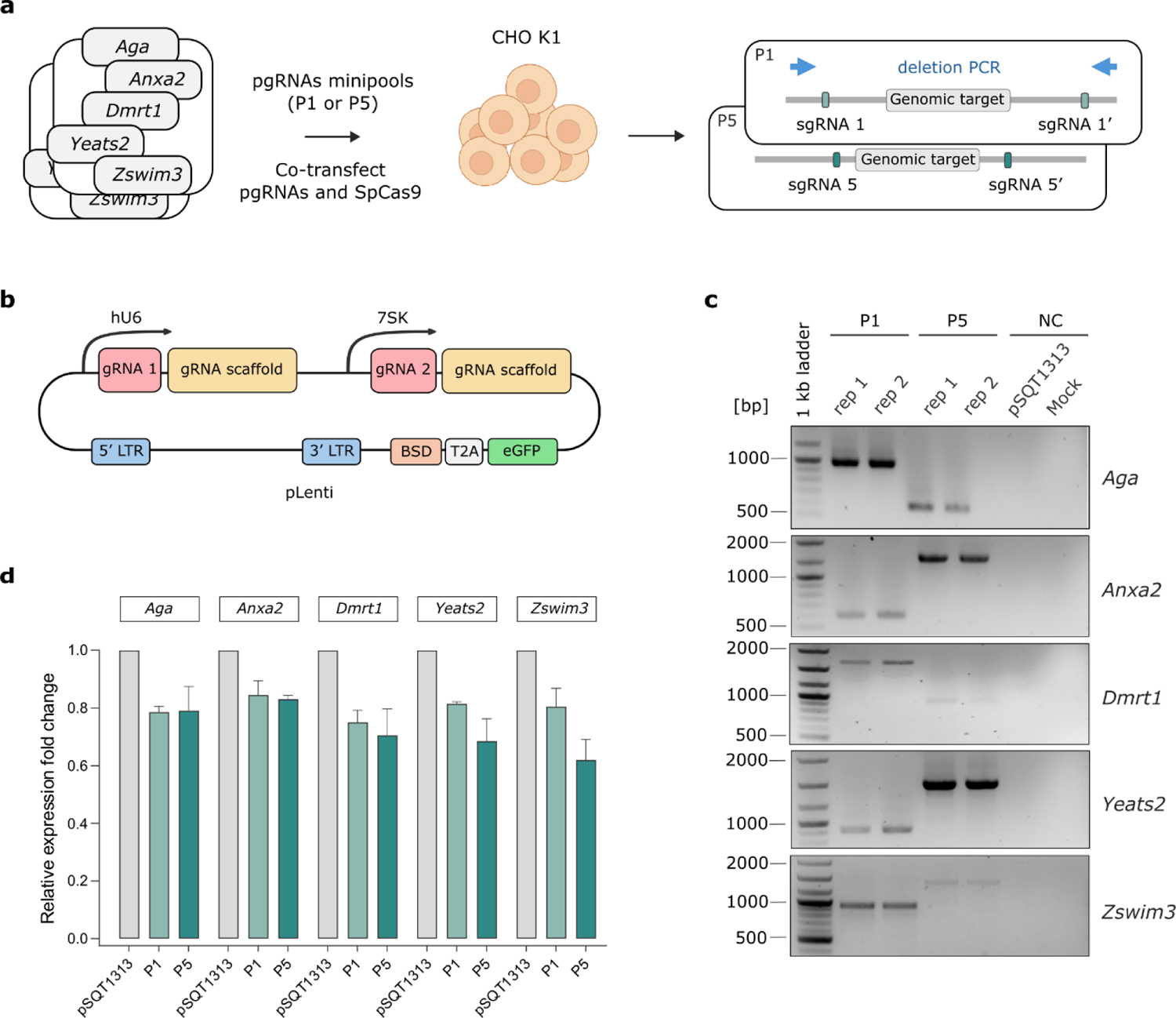
Multiple paired gRNAs delivered in parallel to CHO cells create large deletions at their target genes. (a) Five paired gRNA combinations were designed for each of the five selected genes (Aga, Anxa2, Dmrt1, Yeats2, Zswim3). Two individual pgRNAs pools were created by combining all the gene-specific pairs with the highest (P1) and lowest (P5) Doench scores. The pools were separately and transiently transfected into CHO cells with a SpCas9-expressing plasmid. The guide pairs are designed to introduce strand breaks upstream and downstream of the coding sequence, resulting in the complete deletion of the intervening genomic fragment. Hence, each pgRNA pool simultaneously targeted the five selected non-essential genes (non-EGs), inducing kilobase-scale deletions (12−106 kb). (b) Structure of the plasmid expressing the paired gRNA library (pgRNAs). The U6 promoter (hU6) and both gRNA coding sequences were cloned into a pLL3.7 lentiviral backbone, containing the blasticidin resistance (BSD) gene and eGFP marker along with the sequences required for lentiviral transduction (5’ and 3’ LTRs). Amplified DNA fragments encoding the scaffold for the first gRNA (gRNA 1) and 7SK promoter (7SK) were then cloned in front of the second gRNA (gRNA 2) using the Golden Gate method. (c) Qualitative analysis of the pgRNA-mediated deletion strategy for target genes via deletion PCR. Deletion PCR primers were designed upstream and downstream of the guide-targeting sites. Amplicons were observed only when a successful deletion occurred. Positive bands can be seen for both P1 and P5 while the controls (NC – negative control and untransfected mock) showed no bands, as expected. Deletion PCRs were performed in biological replicates (n=2) (d) Quantitative analysis of the pgRNA-mediated deletion strategy for target genes. Target gene mRNA levels were assayed by qRT-PCR following RNA isolation and cDNA synthesis. Bar graphs show fold changes in mRNA expression, normalized against GAPDH and relative to the negative control (co-transfected with a SpCas9 expressing plasmid and the empty parental plasmid). Data are presented as mean ± SD, n = 3.

Experimental design included untransfected control cells, cells transfected with the empty vector (pSQT1313) and the two populations of cells transfected with either the P1 or P5 pool. Across all tested genes, expression levels decreased by 10% to 45%, with no preferential deletion based on fragment size. Notably, pgRNAs from each pool achieved comparable reductions in gene expression, despite differences in predicted targeting efficiency.

### 3.2 A custom genome-scale CRISPR library for targeted deletion of large genomic fragments

A genome-scale CRISPR library was designed using the software *multicrispr* (Bhagwat et al., 2020), which was adapted to meet our specific pgRNA requirements. *Multicrispr* was chosen over other available CRISPR tools due its superior efficiency in handling large datasets and its capacity for parallel targeting on a genome-wide scale. Using the PICRH CHO genome assembly (Hilliard et al., 2020), chromosomes were first computationally fragmented into 150 kb windows. The first 10 kb within each chromosome end, which likely correspond to low-complexity or telomeric sequences, were excluded, as were scaffolds smaller than 500 kb. For the 1 kb regions upstream and downstream of each window *multicrispr* predicted all potential N20-NGG CRISPR target sites (Figure 2). The pgRNA library was designed to target the entire genome, irrespective of gene presence. Cut sites within exon sequences were excluded to avoid interference with coding regions, while gRNAs predicted within introns were automatically designed to be at least 50 bp distant from exon start sites. Additional filtering parameters were applied to ensure gRNA integrity by excluding candidate guides containing polyT (TTT) motifs (a Pol III transcriptional stop signal), BsmBI restriction sites, Graf motifs in EMS regions (Graf et al., 2019) (**Suppl. table S2C**), and guides with GC content <35% or >75% (T. Wang et al., 2014). In parallel, on-target scoring (*ontargetmethod=“Doench2016”*) was performed to prioritize gRNAs with high predicted targeting efficiency. The resulting gRNAs were ranked according to their *Doench2016* score (Doench et al., 2016) and any guide with a score <0.5 was filtered out. To minimize potential off-target effects, all candidate spacers with one or more genomic off-target sites (*mismatches=0*) were excluded. For each 150 kb window, the highest-ranked guides upstream and downstream were selected, listing them according to their on-target score. Seven or eight guides per window provided redundancy while maintaining library size (Hart et al., 2017a). The resulting genome-scale library consisted of 112,272 gRNA pairs targeting 14,034 genomic regions (windows), irrespective of the presence of coding sequences (**Suppl. table S2A**). The library design effectively covered ∼88% of the PICRH genome, making it the first pgRNA library for CHO cells with near-full genome coverage. A small percentage corresponded to low complexity regions, such as centromeric or telomeric areas, where guide RNA prediction is computationally not feasible, and thus was excluded from the design process.

**Figure 2.**
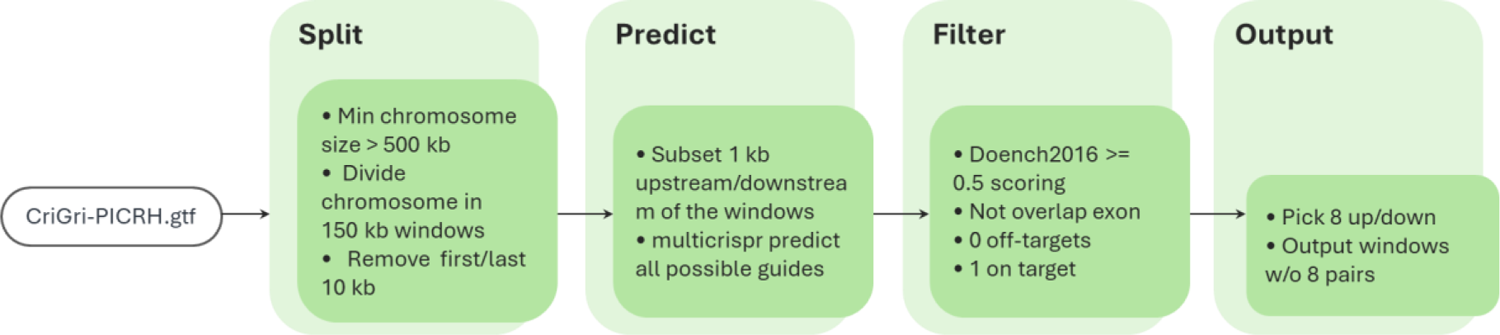
Stepwise pipeline for generating the custom pgRNA genome-scale library

**Figure 3.**
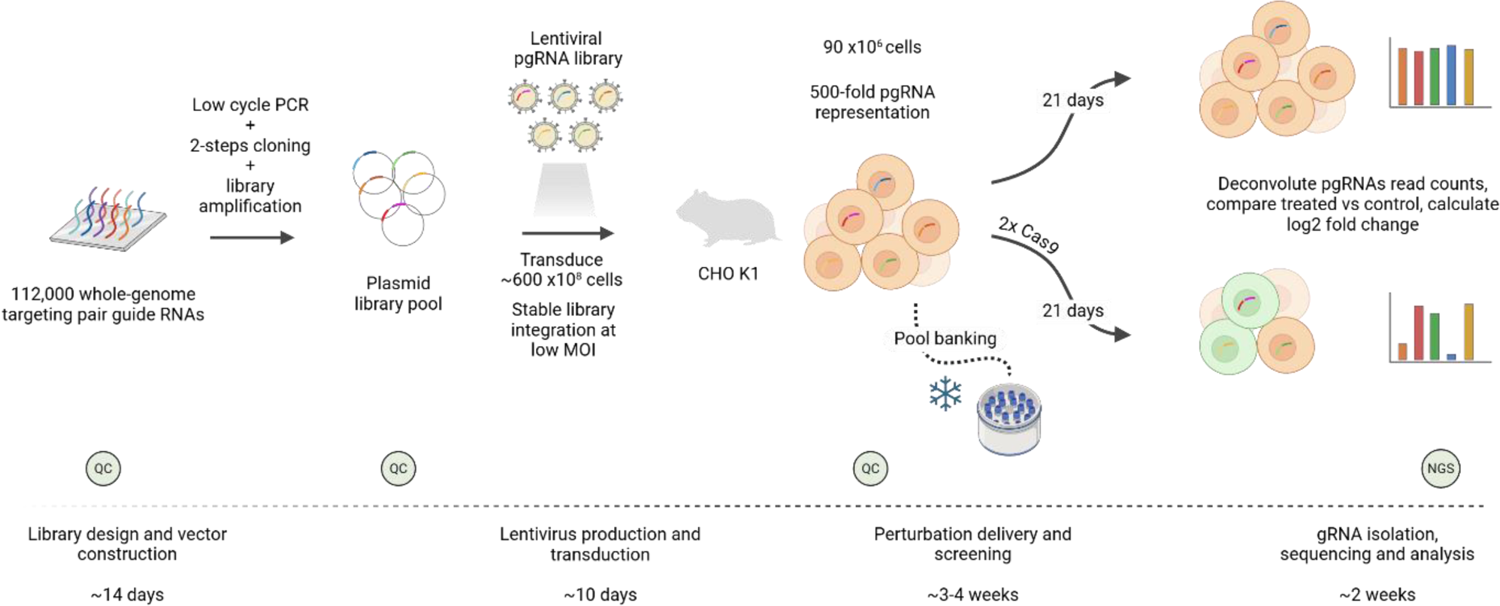
Overview of the dual-guide RNA mediated CRISPR-Cas9 deletion screen. The workflow comprises four major parts: (1) Library design and plasmid library construction, (2) lentiviral library production and transduction, (3) Cas9 perturbation and screening, and (4) next generation sequencing (NGS) of PCR amplified pgRNA pairs. Quality control (QC) samples were collected at three key points throughout the workflow to monitor the distribution of guides and ensure experimental reliability.

### 3.3 Establishment of a rational dropout screening platform for CHO cells

To identify the essential CHO genome, we relied on lessons learned from previous paired guide RNA (pgRNA) deletion screens (Fei et al., 2019; Schmieder et al., 2021; Zhu et al., 2016), refining and optimizing a workflow specifically tailored for our essentiality study. Generating a representative cell pool library requires maintaining a minimum 500-fold coverage per guide pair, which ensures robustness and statistical power for the screen (Bock et al., 2022). In addition, using at least three biological replicates consistently increases the total number of hits identified (Doench et al., 2016; Hart et al., 2017b). To ensure full representation, our 112k pgRNA library was delivered into CHO K1 cells via lentiviral transduction. The protocols were optimized for suspension-adapted cells, improving efficiency and scalability (Napoleone et al., 2025). To minimize a potential source of false positives resulting from integration of multiple library constructs per cell, we targeted a transduction efficiency of 14-16% (MOI ∼0,16), subsequently confirming a vector copy number (VCN) of 1.4 integrations per cell via ddPCR. This ensured that the majority of cells received a single construct, reducing background noise in downstream analyses. Following enrichment for positively transduced cells, >95% of the selected population stably retained the pgRNA library for up to six months without selective pressure (**Suppl.** figure 4), providing a robust and ready-to-go CHO cell platform for essentiality screening. To maintain precise control of perturbation events and avoid spurious editing, SpCas9 was delivered transiently to the cell population. Consistent with findings from (Xiong et al., 2021), a brief SpCas9 enrichment improved editing efficiency. This effect was further amplified by a second perturbation eight days after the first one, significantly increasing the likelihood of CRISPR-induced deletions (**Suppl.** figure 5). To capture the guide distribution across the entire screen, gRNA sequences were PCR-amplified from genomic DNA harvested from the cells at 0 and 21 days post perturbation. These amplicons were analyzed by Illumina sequencing and changes in pgRNA abundance were computed.

### 3.4 Genome-scale dropout screen reveals essential genomic regions

Large-scale pooled screens rely on sequencing pgRNA amplicons as the primary readout. Thus, raw sequencing reads were mapped to the reference guide library, and the relative abundance of each pgRNA was quantified based on read counts. pgRNAs targeting essential genomic regions were expected to show reduction in their read counts, reflecting their depletion in the cell population due to death of compromised cells. In contrast, pgRNAs targeting non-essential regions were anticipated to maintain stable read counts. To generate individual count tables, MAGeCK (Li et al., 2014) and Bowtie2 (Langmead & Salzberg, 2012) were both used to map pgRNA reads to the reference library. The respective count tables were subsequently analyzed using the MAGeCK αRRA algorithm to systematically evaluate individual guide abundance (Figure 4) and perform window-based ranking of altered pgRNA distributions relative to the control (**Suppl.** figure 6). In the absence of additional selective pressures, the SpCas9-mediated perturbation was the sole factor responsible for these deviations. Evaluation of covariance using Pearson correlation demonstrated high homogeneity within the control group (r = 0.92 – 1.00) and greater diversification in the perturbed population (r = 0.76 – 0.78) (**Suppl.** figure 6). Principal component analysis further revealed distinct clustering (**Suppl.** figure 6), with 65% of the observed variation attributed to differences between perturbed and control cells. Overall, the perturbation of pgRNAs distribution confirmed successful depletion of targeted genomic regions. By aggregating fold-change data (**Suppl. table S2B**) across all pgRNA-targeted windows, we identified 427 genomic regions (64 Mb) (Figure 4e) that were significantly depleted across five biologically independent parallel screens, suggesting that the deletion of these 150 kb genomic segments regions impaired cell growth and proliferation of suspension-adapted CHO cells. Of the 427 identified essential regions, 285 (67%) contained multiple genes, either entirely contained within the targeted window, or partially shared among adjacent regions; 81 regions (19%) contained only a single gene within the 150 kb window, suggesting that these genes might be individually essential when disrupted. In contrast, 61 (14%) of the essential windows lacked any gene annotation and thus can be defined as genomic “dark matter” (Figure 4b).

**Figure 4.**
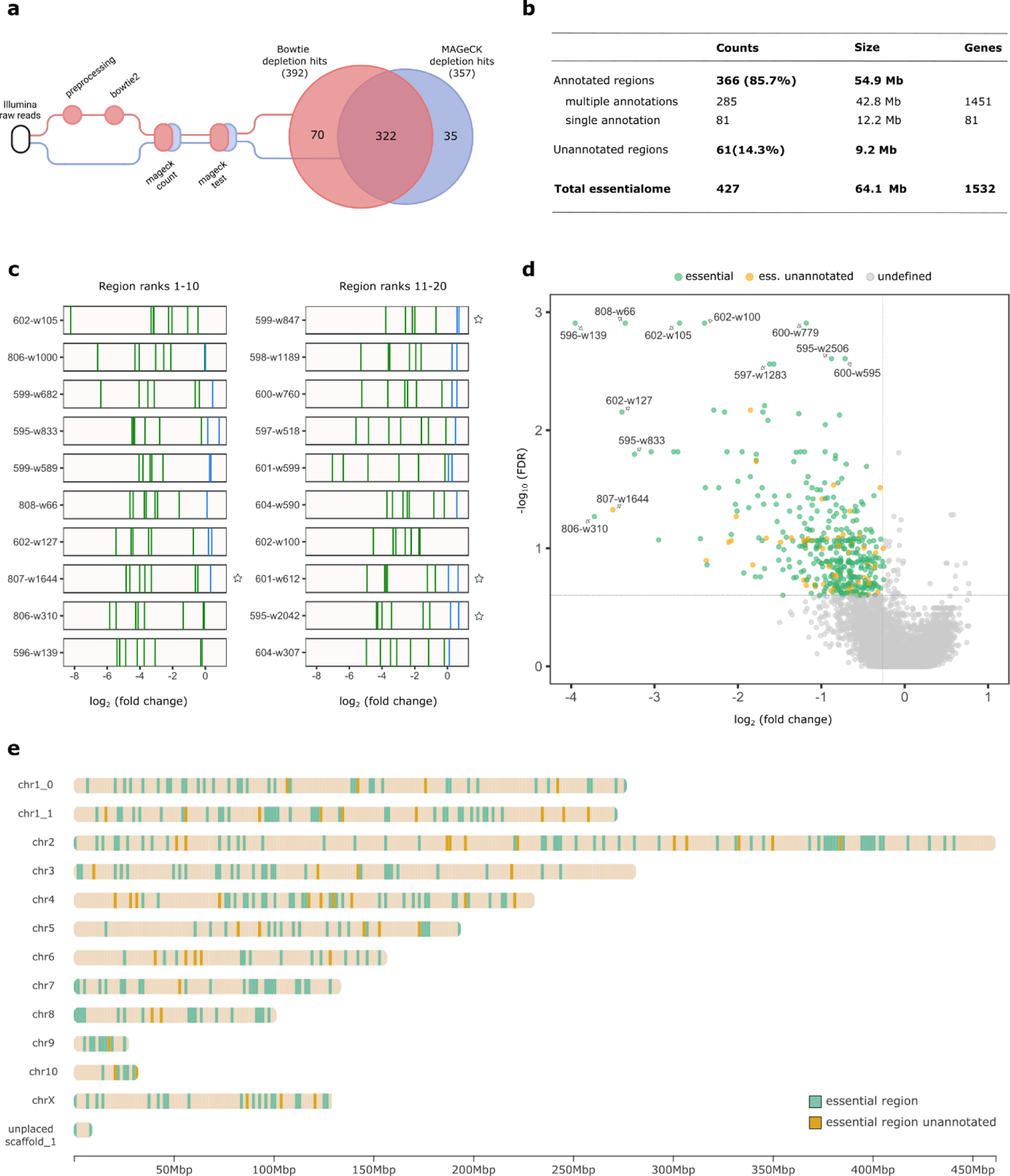
Essentiality dropout screen reveals essential genome regions in CHO cells. (a) Venn diagram of the overlap between the hits identified by mageck test command, using the individual count tables generated by both MAGeCK and Bowtie alignment. (b) Composition of the essential region dataset. (c) The distribution of log2(fold change) of individual gRNA (blue bar: depleted; red bar: enriched) for the 20 most depleted regions as identified by MAGeCK. Depleted regions lacking annotation are indicated by a star symbol. MAGeCK P-values per region are found in **Suppl. table S2**. (d) Volcano plot for the essential hits obtained in the depletion screen. The x-axis denotes the log2fold change of the region-level fold change (median log2(fold change) for all pgRNAs per region. The FDR values in y-axis were derived from the MAGeCK score. Essential regions are differentiated based on the presence of genomic annotation (green) or its absence (gold). (e) Genome-scale distribution of the essential regions across all CHO K1 chromosomes, including unplaced scaffolds. Each coloured bar (green or gold) represents an essential genomic region of 150 kb.

### 3.5 Analysis of the CHO cell essentialome

To profile the biological context of essential regions, we performed over-representation analysis of GO terms for genes included in essential genomic regions. The enrichment analysis was conducted for all three ontologies Biological Process, Molecular Function and Cellular Component (background genes n = 4,510 for biological process, n = 5,283 for molecular function, and n = 5,430 for cellular component). No enrichment of biological process and molecular function terms was found (Figure 5a). Genes in essential regions were enriched in cellular component terms relating to intracellular protein-containing complexes, such as catalytic complex (adjusted *p* = 0.0018), and more specifically histone acetyltransferase complex (adjusted *p* = 0.046). For regions without annotated genes, essentiality could be explained by the presence of elusive transcripts or specific epigenetic features. First, to investigate transcriptional activity within the essentialome, we integrated an RNA-seq dataset (Weinguny et al., 2020) derived from the same cell line with the identified essential regions. The analysis revealed that the lack of annotation for 61 essential regions aligned with the absence of significant transcripts (Riedl et al, 2025). In contrast, the remaining 366 essential regions displayed diverse transcriptional activity profiles, containing both gene expression (260 regions) and lack thereof (106 regions). Next, we assessed the epigenetic context of the essentialome using chromatin states. We used the ChIP-seq dataset published by Feichtinger (Feichtinger et al., 2016), covering six histone marks (H3K4me3, H3K27ac, H3K4me1, H3K36me3, H3K9me3 and H3K27me3), and determined eleven chromatin states based on the latest assembly (Hilliard et al., 2020). This chromatin state annotation was segmented into bins of 200 bp and filtered to contain only segments fully contained within genomic regions covered in our screen. Images of all essential regions, including annotation, chromatin state and RNA-seq read mappings are presented in **Suppl. Materials**. Overlap enrichment of chromatin states in essential genomic regions unveiled significant shifts in chromatin state composition (Figure 5c-e). All but two chromatin states showed significant enrichment or depletion. The largest difference was seen for the Repressed Heterochromatin state defined by the H3K9me3 histone mark, with a fold change of 4.13 over the background (*p* < 0.01) and for the Polycomb repressed state characterized by H3K27me3.

**Figure 5.**
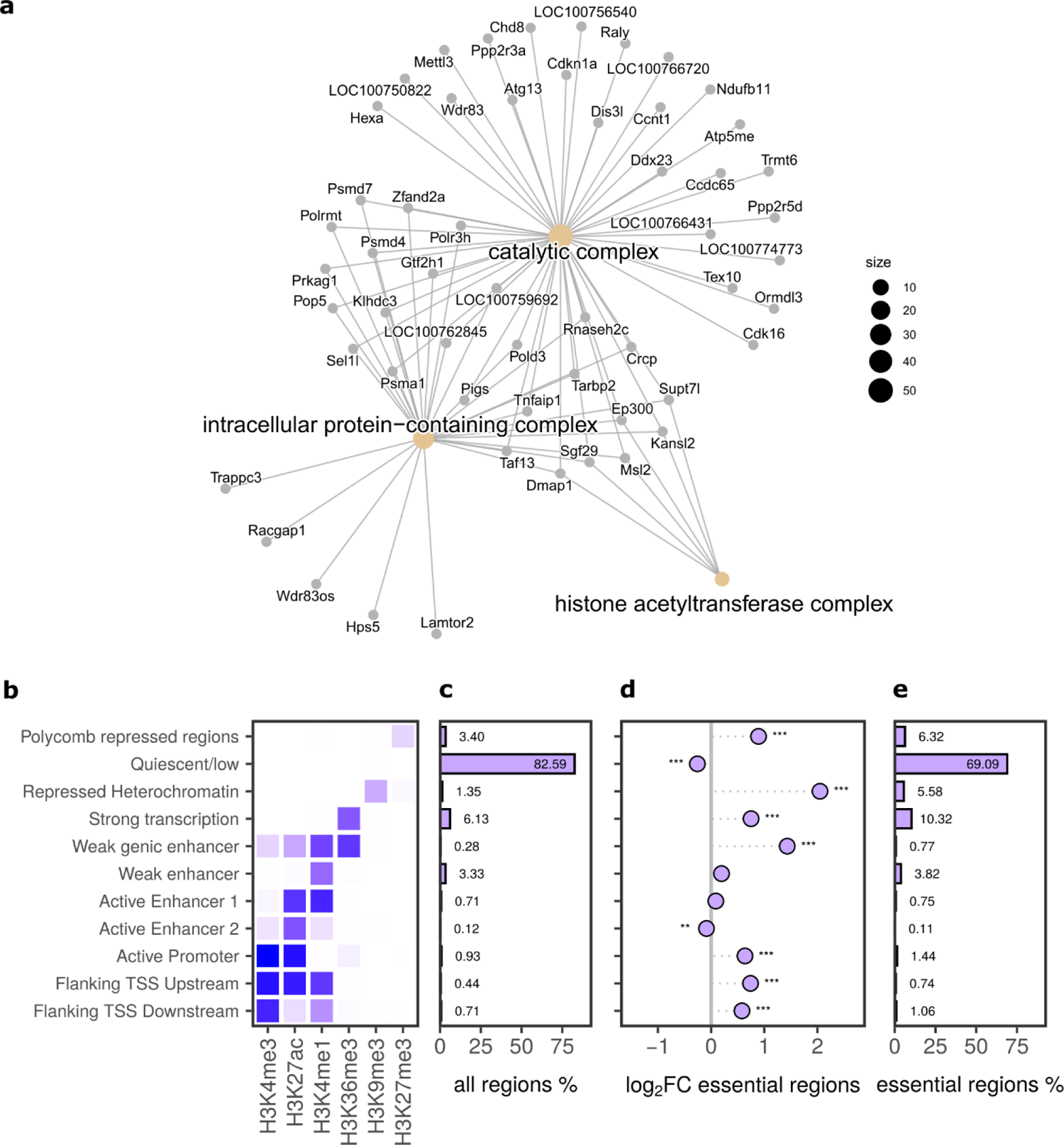
Functional and epigenetic profiling of the CHO cell essentialome. (a) Concept-network plot of enriched GO cellular component terms among genes in essential regions. Evaluated by Fisher’s exact test with Benjamini-Hochberg correction. GO terms were considered enriched at an adjusted p < 0.05. (b) Patterns of six histone marks that define eleven chromatin states. (c) Percent distribution of chromatin states in all regions of the genome. (d) log_2_ fold change of chromatin state distribution in essential regions compared to all regions. Testing for significant enrichment by 2000 permutations resampling from all genomic regions followed by multiple testing correction with Benjamini-Hochberg. (e) Percent distribution of chromatin states in the essential regions identified.

Notably, for the 64 essential regions with annotation, but lacking transcription, these two chromatin states were the most frequently found ones. Also, of the 61 regions without annotation that we called “dark matter”, 24 exhibit no transcription, but contain a chromatin state, predominantly Polycomb repression – despite the absence of any known gene to be repressed. However, 37 regions do not contain any of the histone marks used here, nor any transcripts, so could be called “black matter”.

### 3.6 Validation and functional assessment of essentiality by individual knockout

To verify the results of the screen, six genomic regions predicted to be essential for cell survival were individually targeted for CRISPR-SpCas9 deletion (**Table 1**). We selected two essential regions (ID: 599-w968 and 600-w83) containing coding sequences; the top two regions with FDR < 0.05 characterized by quiescent chromatin and lacking coding sequences (ID: 595-w1249 and 807-w1644); and two regions with the highest proportion of enhancer chromatin marks but devoid of coding sequences (ID: 599-w370 and 601-w290). The two most depleted pgRNAs for each region were individually cloned and co-transfected into CHO K1 cells along with an SpCas9-expressing plasmid. The ability to induce CRISPR-based genomic cuts was first assessed via deletion PCR, where the presence of an amplicon indicated removal of the targeted fragment. Given the essential role of these regions in cell proliferation, we anticipated a gradual reduction in amplicon intensity over time (**Suppl.** figure 7). Further verification of the impact of these deletions was performed using flow cytometry to measure caspase-3/7 activity, a marker of apoptosis, 24 hours after pgRNA-induced excision. All six genomic deletions showed noticeable levels of apoptosis (29 – 45%) compared to cells with a non-essential region deleted or a non-targeting control (Figure 6a). A hallmark of apoptosis is nuclear fragmentation, driven by caspase-3-mediated cleavage of the inner nuclear lamina (Gheyas & Menko, 2023). To confirm the morphological changes induced by deletion of essential regions, we visualized the subcellular apoptotic events using Hoechst and Lamin-B1 immunostaining. These revealed pronounced variations in nuclear architecture depending on the targeted region. (Figure 6b). Nuclear circularity, quantified using Lamin-B1 staining (Matias et al., 2022), has a maximum value of 1, which decreases as nuclear shapes become increasingly convoluted. We observed a significant reduction in circularity in cells with deletions of essential regions, regardless of the presence of coding sequences, indicating increased nuclear deformation (Figure 6c). Together, these results confirmed the efficacy of the screening process in CHO cells, as individual deletions of large genomic regions consistently induced apoptosis. This effect was supported by morphological changes, that led to increased nuclear deformation and decreased nuclear circularity, targeting regions containing both coding and non-coding sequences.

**Figure 6.**
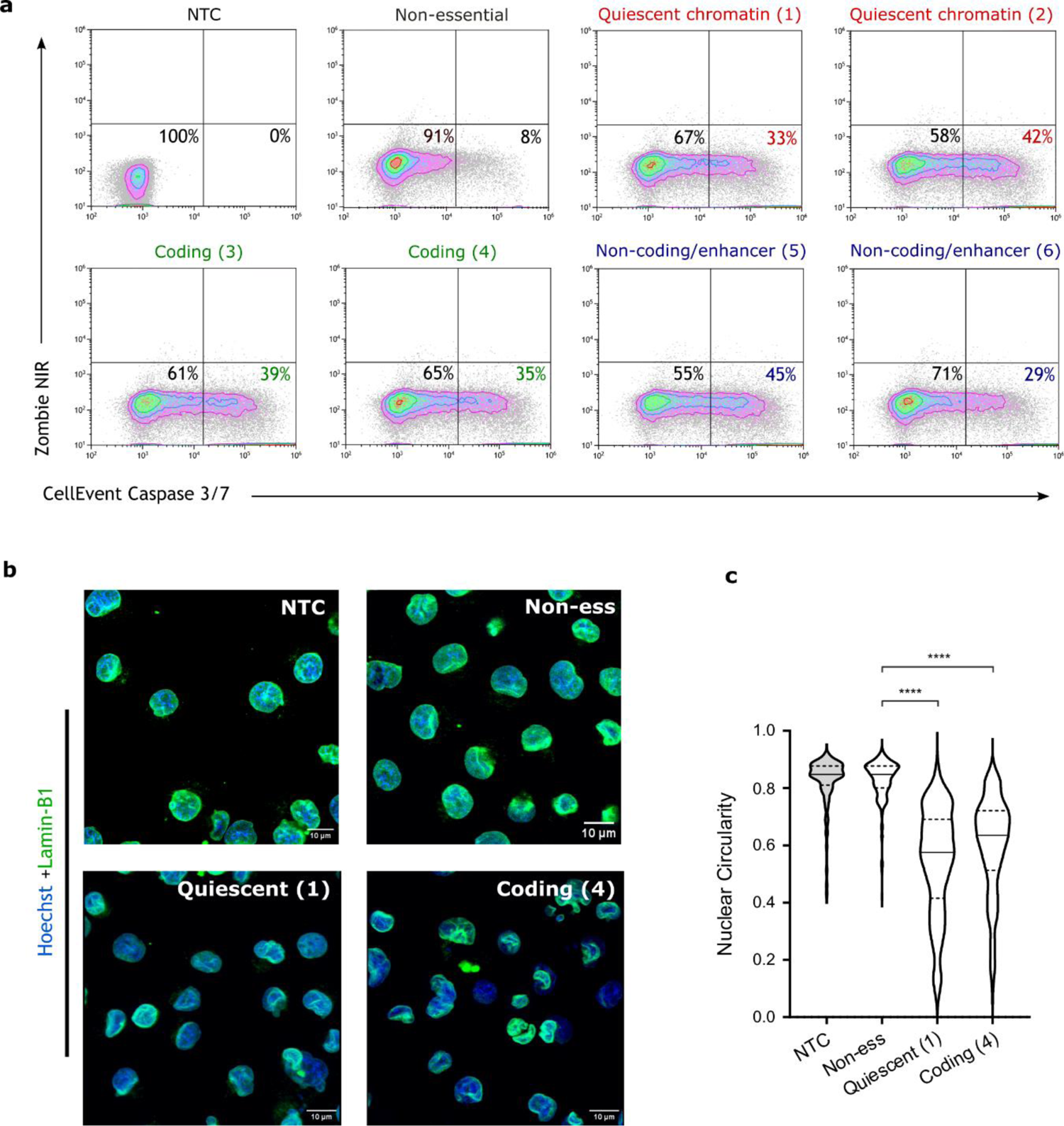
Increased apoptosis and nuclear deformation are associated with the removal of specific fragments of the genome. (a) The increase in the activation of the caspase-3/7 demonstrated a specific apoptosis initiation for each of the six regions selected from the screen. The CellEvent Caspase-3/7 protocol (Invitrogen) was used 24 hours post-transfection with pgRNA and SpCas9. (b) Fluorescence microscopy images of distinct nuclear morphology: regular nuclear structure was maintained in the NTC and non-essential samples, whereas pronounced nuclear aberrations and invaginations were evident in the “coding (4A)” and non-coding (1A)” samples, as shown by lamin-B1 staining. (c) Apoptotic CHO cells exhibited reduced nuclear circularity compared to the deletion of a non-essential region (p < 0.0001).

**Table 1.** List of essential genomic regions selected for individual validation.

| ID | Chromosome | LFC | FDR | Category |
| --- | --- | --- | --- | --- |
| 595-w1249 | NC_048595.1 | -9.92E-01 | 0.038176 | Quiescent chromatin (1) |
| 807-w1644 | NW_023276807.1 | -3.50E+00 | 0.047137 | Quiescent chromatin (2) |
| 599-w968 | NC_048599.1 | -9.52E-01 | 0.03778 | Coding (3) |
| 600-w83 | NC_048600.1 | -5.00E-01 | 0.125645 | Coding (4) |
| 599-w370 | NC_048599.1 | -4.42E-01 | 0.136214 | Non-cod/high enhancer (5) |
| 601-w290 | NC_048601.1 | -4.40E-01 | 0.182226 | Non-cod/high enhancer (6) |

## 4 Discussion

The availability of large genomic datasets and advanced CRISPR screening tools has enabled genome-wide studies of the genotype/phenotype relationship in mammalian cells. So far, most of these studies targeted coding genome regions using the traditional frameshift mutation approach. However, functional analyses of non-coding regions cannot be tackled by this strategy as the impact of sequence changes on folding and structure of RNA is poorly understood (if there is a transcript) (Reber et al., 2018), while even less is known about sequence alterations and their impact on regulatory regions of the genome, such as enhancers or silencers. An alternative screening strategy for these regions is genomic deletion. This approach effectively correlates function with phenotype, regardless of transcription status. By completely removing the region, any function it carries—coding, non-coding, regulatory or currently unknown—is lost, thereby revealing its role (Schmieder et al., 2018). The method has the additional advantage that it is efficient and easily verified by two simple PCRs – a deletion and a non-deletion PCR. In this study, we established a pooled CRISPR screening approach using paired guide RNAs targeted to delete large fragments of the genome (up to 150 kb) to examine the essentiality of the entire genome of CHO cells, irrespective of whether this essentiality is defined by a coding gene or other function. Based on our previous experience with CRISPR screens (Schmieder et al., 2021), a number of potentially critical steps were addressed in order to increase the performance of a genome-scale screen. First, based on data from literature focusing on the efficiency and performance in previously published library screens, several decisions were made on the details of the experimental design. The use of AsCpf1 is already reported in literature to allow fast forward cloning and design of a bicistronic transcription system (Schmieder et al., 2018). However, the more established SpCas9 is often reported to be more active in mammalian cell gene knockout experiments compared to AsCpf1, which failed to show satisfactory separation of essential vs non-essential genes (Liu et al., 2019). The use of dual U6/U6 promoter combination for the expression paired gRNAs (Zhu et al., 2016) was reported to be prone to recombination within promoter sequences in HEK293 cells (Gasperini et al., 2017). Therefore, and since no data was available for CHO cells, we decided to use SpCas9 rather than the previously used AsCpF1, to switch to two different promoters for the two guides in each pair (U6/7SK) and to implement an optimized gRNA scaffold sequence as described by (Dang et al., 2015). Second, the guide library was designed to accommodate the large scale of a CRISPR screen for an entire mammalian genome. As many as 101 software tools for designing gRNAs are described in literature (Torres-Perez et al., 2019). However, as reported by (Bhagwat et al., 2020) only seven can perform off-target analysis and on-target scoring (CHOPCHOP, CRISPOR, CCTop, CRISPRseek, CLD, CRISPRETa and FlashFry). The problem with all these design tools is that they are not developed for a multi-targeting tiling array-like approach required to design a library on a genome-scale target set such as the CHO cell genome. To overcome this obstacle, we developed an R script tool to work with *multicrispr* (Bhagwat et al., 2020), a software that exhibits a much-reduced computing time compared to others and allowed us to accommodate the addition of custom parameters directly correlated to the efficiency of the guides. Thus, we successfully generated a pgRNA library that covers over 88% of the PICRH genome consisting of 112,272 unique pairs of guide RNAs, only excluding low complexity sequences and small scaffolds. Third, in practical terms, we ensured the high quality of the library i) by using high proof-reading capacity polymerase, ii) by the low number of cycles for PCR and iii) by performing NGS for all the preliminary steps before the generation of the actual library cell pool, which is often overlooked for genetic screens. For pooled library screens, the lentivirus-based delivery of library components is the most commonly used system (Piccioni et al., 2018; T. Wang et al., 2014). However, CHO cells growing in suspension are also less easily transduced than other mammalian cell lines, and lentiviruses may elicit unexpected changes in the cellular phenotype (Sack et al., 2016). In contrast, the integration of a guide library by recombinase mediated cassette exchange (RMCE) is known to be a valid alternative, as it is known to create a more controlled system (Baumann et al., 2017; Xiong et al., 2021). However, establishing a landing pad host cell line requires pre-engineering and thus may lead to inflexibility in the choice of target cell line (Pristovšek et al., 2019; Sergeeva et al., 2020).

Moreover, in the context of a genome-scale library, to maintain a redundancy of at least 500-fold in the distribution of guide RNAs across the pool, careful consideration must be given to the efficiency of the delivery method. In our screen, lentiviral transduction achieved an efficiency of up to 20% after careful optimisation of all steps, while maintaining a high library coverage (Napoleone et al., 2025). This considerably reduces the burden of transfecting a large number of cells and makes selection of positive transfectants easier.

Additionally, we have also proven that excisions using paired sgRNAs can be multiplexed with good efficiency. Although a larger deletion in CHO cells of up to 864 kb has also been recently reported (Jerabek et al., 2024), this approach targeted only a specific cluster of non-essential genes. Smaller deletions, such as the 150 kb excisions used in our study, offer a finer and more precise approach to interrogate the genome, allowing a detailed resolution. In our work, while screening for the essentiality of the entire genome, we hypothesize that genomic regions not affected by perturbations, represent areas where genomic excision, does not hinder the cell proliferation. These unaffected portions can be described as non-essential for cell survival, opening up to the potential generation of a set of regions that could simultaneously be excised from the CHO cells to reduce its genome.

In this study, we defined an essential region as any genomic region of a defined length in the CHO genome whose deletion decreases cell growth and proliferation. We demonstrated that 427 areas of the CHO genome fall into this category, irrespective of the presence or absence of coding or non-coding sequences. By doing so, we revealed for the first time that genomic regions can be essential despite the fact that the genes present in this sector are not transcribed, or in the complete absence of any known annotated gene or chromatin state functionality. If silenced genes are present in a given region, one could argue that they may not be transcribed at the time of sampling for RNAseq, but that they may be required in stress situations currently not encountered by the cell, where the absence of such a gene then would entail cell death. Remarkably, the majority of not-annotated regions that were classified essential and carried a chromatin state mark were characterized by polycomb repression – a poised state of chromatin where cells keep a gene inactive but ready for re-activation (Macrae et al, 2023). This would indicate that these regions encode for a reserve function only needed under certain circumstances, but currently not known or annotated. Finally, for the 37 regions that did not even contain a known chromatin state, their essential function is a black box. In general, accounting for a CHO genome size of 2.45 Gb (Xu et al., 2011) and a deletion resolution of 150 kb, we propose that only 2-5% of the CHO genome is indeed essential for cell survival.

There are some potential limitations to the current study. Biases in CRISPR screens are particularly due to differential gRNA efficiency across targeted regions. These biases may skew the results, necessitating repeated studies to fine-tune the efficiency of the guide library. Computational methods are per se a “best guess” system, where each guide pair has multiple factors that can limit their functionality. Among these, chromatin accessibility is a major determinant of binding efficiency (X. Wu et al., 2014), with potential implications for affecting the outcome of SpCas9 perturbation-based screening. A potential approach could be sub-setting the original library and focusing on additional screens only for the regions that did not experience any initial dropout. A second aspect to consider is that the identification of essential regions is highly context-dependent. The results obtained in our screen are valid for CHO K1 cells, which are a non-producer cell line cultivated under standard conditions. Other CRISPR studies have already characterized genes responsive to specific stressing cultivation conditions (S. H. Kim et al., 2023) and similar approaches could be reproduced with essentiality as a target. It can be assumed that a core set of essential regions shared across mammalian cell lines and valid for most cultivation conditions exists. However, due to the incompleteness of each screen, multiple essentiality screens are required in order to account for all relevant conditions and genomic regions.

In summary, our findings demonstrated the feasibility of screening for essentiality in a pooled format using a paired gRNA approach. Here, we presented a list of essential genomic areas that must be carefully considered when applying genomic engineering strategies in CHO cells. We provided a valuable strategy to screen for the essentiality of the non-coding genome of mammalian cells, obtaining a fascinating peek inside an elusive, but large portion of the CHO genome, which has frequently been neglected. The result, on one hand, highlights how far we are from a full functional understanding of the entire genome; on the other hand, it emphasizes how non-coding regions may develop into an appealing target towards a more rational cell line development.

## CRediT authorship contribution

**Federico De Marco:** Investigation, Methodology, Formal analysis, Validation, Visualization, Writing - original draft, Writing - review & editing. **Ivy Rose Sebastian:** Investigation, Methodology, Validation, Writing - review & editing. **Antonino Napoleone:** Investigation, Methodology, Writing - review & editing. **Alexander Molin:** Formal analysis, Data curation, Visualization. **Markus Riedl:** Software, Formal analysis, Resources, Visualization. **Nina Bydlinski:** Conceptualization, Methodology. **Krishna Motheramgari:** Software, Methodology. **Mohamed K. Hussein:** Supervision, Project administration, Writing - review & editing. **Lovro Kramer**: Resources. **Thomas Kelly:** Resources, Supervision. **Thomas Jostock**: Resources, Supervision. **Nicole Borth:** Conceptualization, Funding acquisition, Supervision, Writing - review & editing.

## Declaration of competing interests

Lovro Kramer and Thomas Jostock are employees of Novartis. Thomas Kelly is an employee of Johnson & Johnson Innovative Medicine.

## Acknowledgements

We thank the entire Borth laboratory for support and advice, in particular Ursula Kiesswetter and Zerina Arnautalic for excellent lab organisation. We thank the BOKU Core Facility Biomolecular & Cellular Analysis (BmCA) and the Next Generation Sequencing Facility at Vienna BioCenter Core Facilities (VBCF), member of the Vienna BioCenter (VBC), Austria. This work was supported by the Austrian Center of Industrial Biotechnology (acib): the COMET center acib: Next Generation Bioproduction is funded by BMK, BMAW, SFG, Standortagentur Tirol, Government of Lower Austria and Vienna Business Agency in the framework of COMET - Competence Centers for Excellent Technologies. The COMET-Funding Program is managed by the Austrian Research Promotion Agency FFG. Ivy Rose Sebastian and Markus Riedl appreciate support by the FWF PhD Program Grant # W1224 “Biotechnology of Proteins – BioTop”.

## Abbreviations

Aga: Aspartylglucosaminidase
Anxa2: Annexin A2
CHO: Chinese hamster ovary
CRISPR: Clustered Regularly Interspaced Short Palindromic Repeats
ddPCR: droplet digital PCR
Dmrt1: Doublesex And Mab-3 Related Transcription Factor 1
eGFP: enhanced green fluorescent protein
FDR: false discovery rate
Gapdh: Glyceraldehyde 3-phosphate dehydrogenase
GO: gene ontology
gRNA: guide RNA
HEK293: human embryonic kidney 293 cells
MAGecK: Model-based Analysis of Genome-wide CRISPR/Cas9 Knockout
MOI: multiplicity of infection
PCR: polymerase chain reaction
pgRNA: paired gRNA
qRT-PCR: quantitative reverse transcription PCR
sgRNA: single gRNA
SpCas9: Streptococcus pyogenes Cas9
VCD: viable cell density
VCN: vector copy number
Yeats2: YEATS Domain Containing 2
Zswim3: Zinc Finger SWIM-Type Containing 3

